# Applying taxonomic boundaries for species identification (ABIapp): A convenient and accurate application for species delimitation of parasitic helminths

**DOI:** 10.1101/2022.06.15.496221

**Authors:** Abigail Hui En Chan, Urusa Thaenkham, Tanaphum Wichaita, Sompob Saralamba

**Author notes:** Email addresses: AC UT TW SS.

## Abstract

**Background:** Parasitic helminths are highly diverse and ubiquitously distributed in various environments and hosts. Their vast species diversity renders morphological and molecular species delimitation challenging, due to phenotypic and genotypic variations. Currently used approaches to species delimitation are generally computationally intensive. Here, using genetic distances, we developed and validated a simple and easy-to-use application, Applying taxonomic Boundaries for species Identification (ABIapp), to aid in helminth species delimitation.

**Methodology/Principal Findings:** ABIapp uses a database of cut-off genetic distances obtained using the K-means algorithm to determine helminth taxonomic boundaries for ten genetic markers: The nuclear 18S and 28S rRNA genes, ITS1 and 2 regions, and the mitochondrial 12S and 16S rRNA, *COI, COII, cytB*, and *ND1* genes. ABIapp was written in R, and the Shiny framework was used to produce an interactive and user-friendly interface. ABIapp requires just three types of input (genetic distance, genetic marker, helminth group) that are easily generated through basic morphological and molecular analysis. To validate ABIapp’s accuracy and robustness for use, validation was performed both *in silico* and with actual specimens. Prior to validation, ABIapp’s database of genetic distances and species used was increased to broaden the app’s applicability. *In silico* validation was conducted by obtaining 534 genetic distances from 91 publications and inputting these into ABIapp. Using confusion matrices, an overall classification accuracy of 79% was achieved, demonstrating the robustness and accuracy of ABIapp. Using sequences of the 12S, 16S, *COI*, and 18S rRNA genes obtained from ten representative helminth specimens, an overall classification accuracy of 75% was achieved.

**Conclusions/Significance:** Our results demonstrate the applicability and robustness of ABIapp for helminth species delimitation using ten common genetic markers. With a user-friendly interface, coupled with minimal and simple data input and robust classification accuracy, ABIapp provides helminth researchers with a convenient tool for helminth species delimitation.

**Author summary:** Species delimitation of organisms is often an issue of debate, with varying criteria used to determine species boundaries. Helminths are no exception, and their vast species diversity renders species delimitation challenging due to both physical and genetic variations. Moreover, as climate change progresses, helminths are also adapting to the changing environment through morphological and molecular changes. These variations render it challenging for helminthologists to determine whether a particular helminth belongs to the same or a different species. We have developed an application, ABIapp, a simple tool to aid helminth species delimitation using genetic distances; this app is readily available for a wide audience. Encompassing ten genetic markers for the three parasitic helminth groups (nematodes, trematodes, and cestodes), ABIapp uses cut-off genetic distances generated via machine learning to define species boundaries at each taxonomic level. To use ABIapp, just three types of information are needed, requiring only basic morphological and molecular expertise. We validated ABIapp using both mathematically modeled genetic distances and actual specimens and demonstrated a classification accuracy of 79% and 75%, respectively. This new, convenient, and validated application for helminth species delimitation will aid species identification applicable in the fields of helminth taxonomy, disease diagnosis, and biodiversity.

## Introduction

Parasitic helminths, comprising the Nematoda and Platyhelminthes, are highly diverse and globally distributed [1, 2]. Estimates of helminth diversity remain controversial, with many uncertainties due to the small proportion of these organisms so far described. Based on the Host-Parasite Database held by the Natural History Museum in London, Carlson *et al*. (2020) estimated a global total of 100,000 to 350,000 helminth species (including Phylum Acanthocephala), with 85% to 95% of these being unknown [2]. The large species diversity of helminths can be attributed to, among other factors, their complex life cycles; their ability to host-switch, resulting in rapid adaptive radiation; parasitizing a variety of hosts; and their ubiquitous presence in various ecological habitats, such as soil and marine environments [3, 4].

Traditionally, morphological characteristics have been used for helminth species identification. However, accurate species identification using classical taxonomy can be hindered by ambiguous morphological characteristics, phenotypic plasticity due to a variety of hosts and habitats, technical differences in the preparation of specimens, and incomplete specimens that lack key diagnostic morphological characteristics [5-7]. The molecular era gave rise to an alternative tool in the form of genetic markers for species identification. The use of genetic markers not only accelerated the pace of molecular-based identification of helminths but also enabled accurate species identification of previously morphologically indistinguishable species [6-9]. Despite the advantages of molecular-based identification, however, the existence of species complexes and cryptic species complicates its use [2, 8, 9]. It has been estimated that there are on average 2.4 cryptic species per cestode species, 3.1 for trematodes, and 1.2 for nematodes [2]. Genotypic variation renders it challenging to define species boundaries and agree on what constitutes whether a species is the same or different (species delimitation). Generally, when using mitochondrial genes, different species have a 5% to 10% genetic variation between them [7]. However, in assessing ten general genetic markers for parasitic helminths, Chan *et al*. (2021) revealed wide genetic variations among helminths and showed that a general yardstick of genetic distances is not appropriate for helminth species delimitation [10]. The ten genetic markers assessed were the nuclear 18S and 28S ribosomal RNA (rRNA) genes, nuclear internal transcribed spacer 1 and 2 (ITS1 and ITS2) regions, the mitochondrial 12S and 16S rRNA genes, the mitochondrial protein-coding genes of cytochrome *c* oxidase subunits 1 and 2 (*COI* and *COII*), cytochrome b (*cytB*), and NADH dehydrogenase subunit 1 (*ND1*).

Various methods have been used for species delimitation of organisms, including those requiring phylogenetic reconstruction and those relying on distance-based calculations. Among those requiring phylogenetic reconstruction, the Bayesian modeling approach, the General Mixed Yule Coalescent (GMYC) and multi-coalescent model approach, and the Poisson tree processes (PTP) model to infer putative species boundaries are prominent examples [11-15]. Pons *et al*. (2006) used the GMYC model to model speciation in beetles through a neutral coalescent process [11]. Using multiple loci, Yang and Rannala (2014) conducted a simulation study to integrate gene trees [14]. The PTP model, which directly models the speciation rate by using the number of substitutions, was also tested on the *Rivacindela* dataset [15]. Among helminths, the Bayesian species delimitation approach has been applied to identify the cryptic species within *Stegodexamene anguillae*, using mitochondrial and nuclear genetic markers [16]. However, these models are not only computationally intensive but they also require the input of estimated parameters for species delimitation, the prior construction of phylogenetic trees, have a preference for multiple loci, and have not been utilized extensively for helminths. Distance-based approaches for species delimitation include the Automatic Barcode Gap Discovery (ABGD) method, which utilizes the gap between inter- and intra-species variation to partition datasets [17]. The ABGD method has been utilized to delimit the plant-parasitic nematode genus *Pratylenchus*, as well as *Clinostomum* species [18, 19]. However, its use is limited to the mitochondrial *COI* gene. An alternative simple and convenient approach for helminth species delimitation was developed by Chan *et al*. (2021), using the unsupervised machine learning K-means algorithm [10]. The use of K-means genetic distances provides an easy and non-computationally intensive approach for species delimitation.

Here, to make the use of K-means species boundaries for helminths available to a wider audience, we describe the development of a user-friendly application: Applying taxonomic Boundaries for species Identification (ABIapp). With a wide choice of ten genetic markers, users can select the marker that is most appropriate for their research objective. ABIapp provides a convenient and easy-to-use platform for helminth researchers, requiring just simple bioinformatic analysis to aid in species delimitation. To broaden the range of taxa that can be analyzed, we first increased the number of sequences used in the database to update the estimated K-means cut-off genetic distance values for ABIapp. Next, the applicability of ABIapp and its robustness for use were validated by evaluating its species delimitation classification accuracy against genetic distances obtained from previous publications (that used *in silico* methods) and actual specimens. *In silico* validation was performed solely based on an assessment of ABIapp’s classification accuracy and was not used to determine which approach is correct or incorrect. Our work has resulted in a validated, accurate, and convenient application for helminth species delimitation, which is now readily available to helminth researchers.

## Methods

### ABIapp development and workflow

The ABIapp was written in the R programming language, while the Shiny web application framework was used to produce an interactive and user-friendly interface [20]. The source code for ABIapp can be found at https://github.com/slphyx/ABI. ABIapp uses the database of helminth genetic distances to generate species delimitation boundaries by the K-means machine learning algorithm, previously compiled by Chan *et al*. [10].

Designed as a simple and convenient tool for helminth species delimitation, three parameters are required as the inputs for ABIapp. The three parameters are the helminth group of interest, the genetic marker used to generate the genetic distance for the helminth of interest, and the resulting genetic distance value obtained from pairwise sequence analysis. The output shows the results of the queried genetic distance, a graphical visualization, and a table of the genetic distance ranges for the selected marker. The output obtained via ABIapp can reveal 1) the probable classification of the queried genetic distance based on the taxonomic hierarchy level, 2) the range of estimated genetic distances with the selected genetic marker, 3) a probable answer whether the specimen of interest belongs to the same species (based on the species compared during sequence analysis), and 4) suggestions based on the results obtained. Figure 1 shows the ABIapp interface and an example of the results generated.

**Fig 1.**
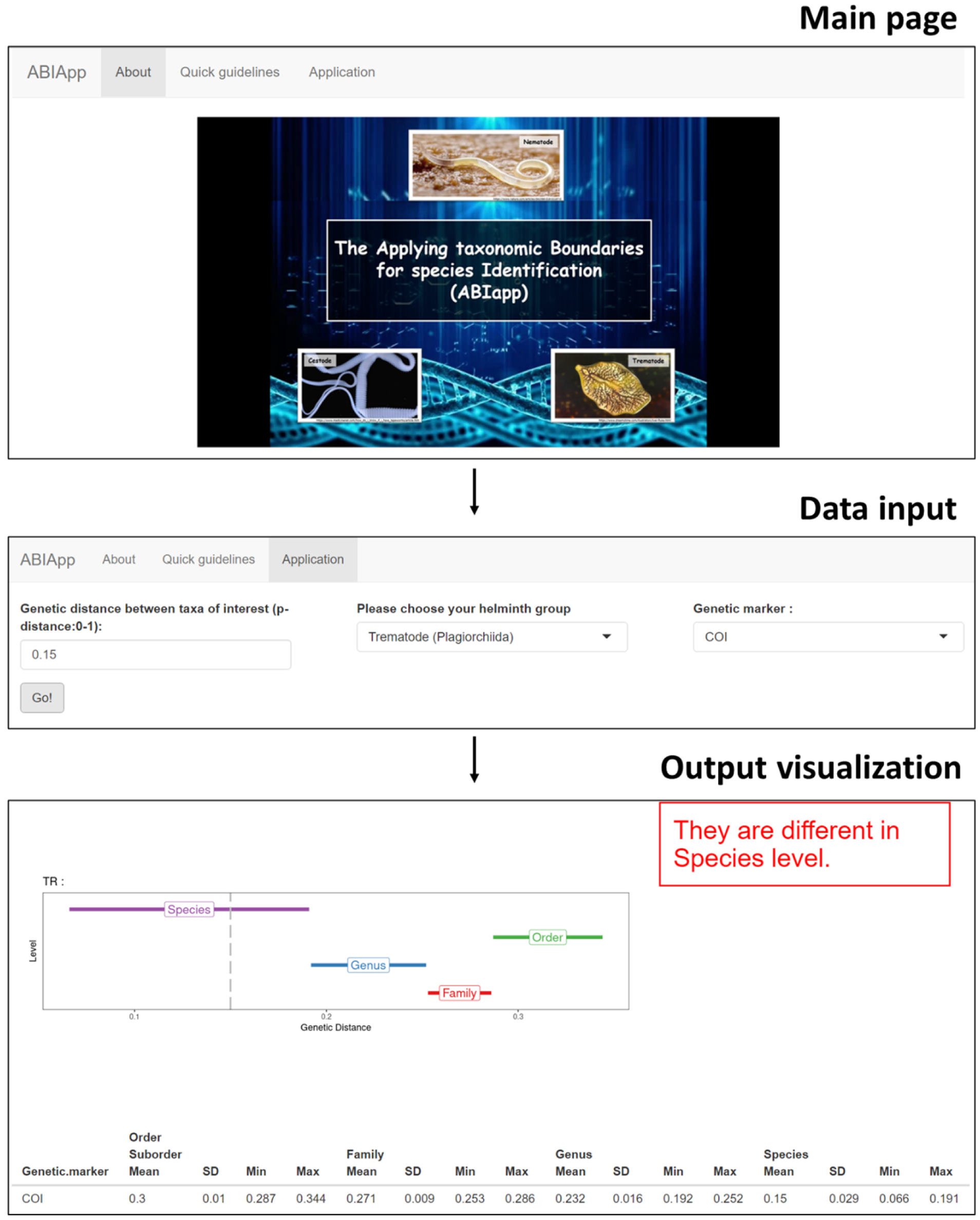
The ABIapp interface showing the main page, types of data to input, and an example of the results generated.

Prior to using ABIapp, morphological identification of the helminth of interest should be performed to obtain some idea of which order or family the queried specimen belongs to. However, in cases where the specimen is unidentifiable, other biological or clinical information may be used to determine the probable helminth group. Next, genetic sequence information must be obtained from one of the ten genetic markers used in ABIapp. The genetic sequence obtained is then compared with genetically closely related species for sequence analysis and to generate pairwise genetic distances for data input into ABIapp.

ABIapp is freely available at https://slphyx.shinyapps.io/ABIApp/ and can also be installed as an R package. A step-by-step guide and more information can be found on the webpage.

### Increasing database taxa and sequences

The database of helminth genetic distances was updated by increasing the number of helminth sequences used to generate the estimated cut-off genetic distance values with K-means. Following Chan *et al*. (2021), helminth sequences of the ten genetic markers (nuclear 18S and 28S rRNA genes, the nuclear ITS1 and ITS2 regions, and the mitochondrial *COI, COII, cytB, ND1*, and 12S and 16S rRNA genes) were mined from the NCBI database [10]. For the mitochondrial genes, full-length sequences were obtained from complete mitochondrial genomes, while full-length or close to full-length sequences were chosen for the nuclear genetic markers. Helminths were also divided into six groups – nematode clade I (Trichocephalida), nematode clade III (Ascaridida and Spirurida), nematode clade V (Strongylida), trematode (Plagiorchiida), trematode (Diplostomida), and cestode. The division follows the taxonomic classification by Blaxter *et al*. (1998) and Olson *et al*. (2003) [21, 22].

Briefly, sequence alignment was performed in ClustalX 2.1 and Bioedit 7.0 for each genetic marker per helminth group [23, 24]. Pairwise genetic distance calculation was then performed in MEGA X to obtain genetic distance values for each genetic marker per taxonomic hierarchy group per helminth group [25]. The genetic distance values obtained were then input into Wolfram Mathematica 12.1 to generate estimated cut-off genetic distance values for each taxonomic level via the unsupervised K-means clustering algorithm [26]. The number of clusters selected was based on the taxonomic levels of the genetic distance values – for example, four clusters represent “species”, “genus”, “family”, and “order”. The helminth sequences used for the database and estimated K-means generated are shown in S1 and S2 Tables, respectively.

### ABIapp validation

#### In silico validation

For the *in silico* validation, available helminth genetic distances were mined from publications and input into ABIapp to determine its accuracy in result prediction. The purpose of obtaining published genetic distances was solely to assess the classification accuracy of ABIapp. The criteria for publication selection were 1) inclusion of genetic distances that were not already included in the database for ABIapp and 2) exclusion of publications without phylogenetic analysis. Taxonomic information and genetic distances were compiled for each genetic marker, helminth group, and which taxonomic hierarchy group the genetic distance obtained belonged to (within species, species, or genus). These taxonomic levels were selected as they are the levels most commonly used in molecular-based studies. We tried as much as possible to obtain genetic distance information for all ten genetic markers and all helminth groups. However, information for some genetic markers was limited due to the scarcity of available publications. A list of publications used is shown in S3 Table.

The compiled genetic distances were then input into ABIapp to determine its accuracy for species delimitation classification. If the genetic distances obtained comprised a range of values, we input the minimum and maximum values. The predictions were classified as either “correct” or “incorrect” (predicted) based on the genetic distance and classification previously obtained based on published data (actual). For example, a “correct” classification at the species level implied that ABIapp correctly predicted the actual genetic distance value at the species level, while an “incorrect” classification implied that ABIapp did not predict the actual genetic distance value at the species level (S4A Table). A confusion matrix was generated based on the predicted results using ABIapp and the actual results for each genetic marker per helminth group, to determine the classification accuracy and error rate [27] (S4B Table). The classification accuracy is calculated as

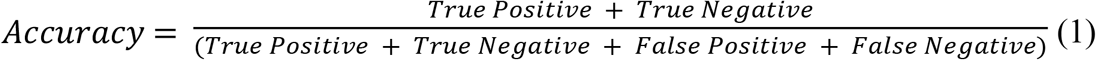

while the error rate is calculated as

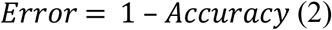

#### Validation with actual specimens

Representative helminth specimens that had previously been archived in the Department of Helminthology, Faculty of Tropical Medicine, Mahidol University, Bangkok, were used to validate ABIapp. Ten specimens comprising representatives of each of the six helminth groups were selected for molecular analysis. These specimens were selected as we were 1) either unable to identify them to the species level or 2) unsure of the accuracy of their morphological species identification. Information about the specimens selected is shown in Table 1.

**Table 1:**
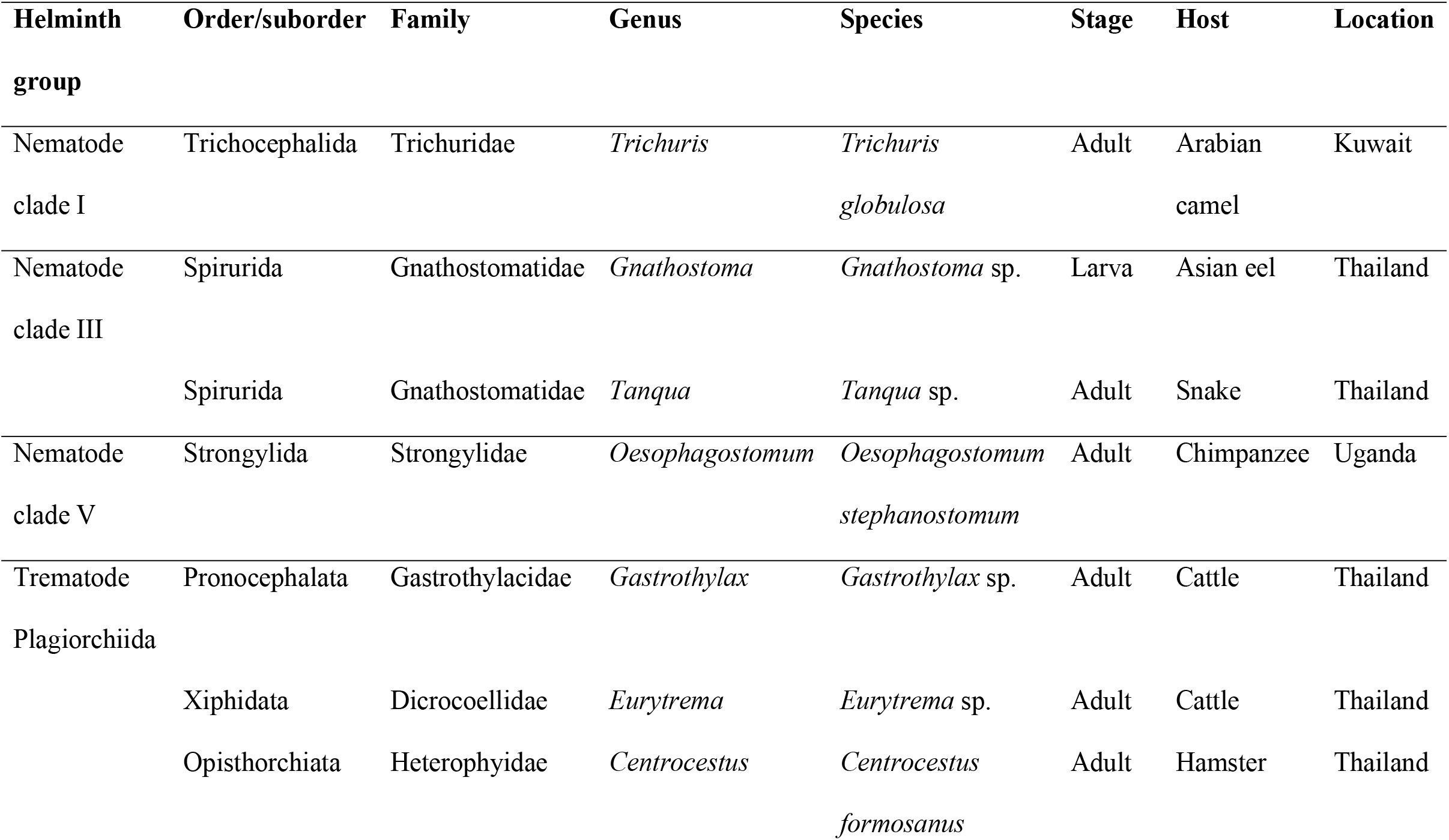
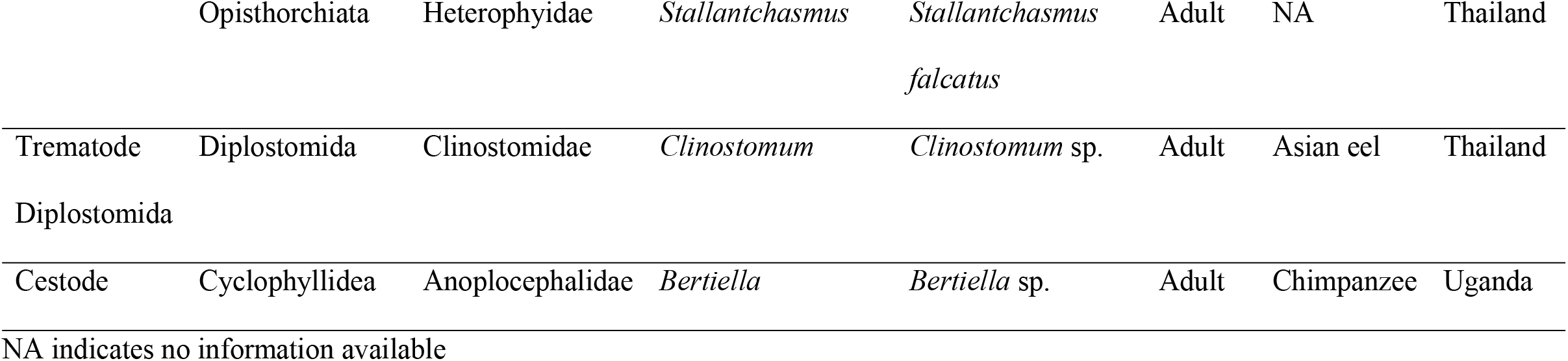
Actual specimens used to validate ABIapp.

For molecular analysis, helminth specimens were individually separated into 1.7-ml microcentrifuge tubes and washed thoroughly with sterile distilled water. A small section of larger-sized specimens was removed for DNA extraction while the remainder of the specimen was stored in 70% ethanol as a voucher specimen. For smaller-sized specimens, the whole specimen was used. Tissue homogenization was performed with silica beads in lysis buffer using a TissueLyser LT (Qiagen, Hilden, Germany). Total genomic DNA was isolated from each sample using the DNeasy® Blood & Tissue kit (Qiagen, Hilden, Germany), according to the manufacturer’s instructions.

We selected the mitochondrial 12S and 16S rRNA, *COI*, and the nuclear 18S rRNA genes as representative genetic markers for molecular analysis. These four genetic markers were selected as there are available primers targeting a broad species range for nematodes, trematodes, and cestodes. Table 2 shows the primers used for each specimen. PCR was performed in a final volume of 30 µl, containing 15 µl of 2X i-TaqTM mastermix (iNtRON Biotechnology, Gyeonggi, South Korea), 10 µM to 50 µM of each primer and template DNA. The thermocycling conditions followed those detailed in the respective publications describing the primers [28-33]. Amplicons were visualized on 1% agarose gel stained with RedSafe® (Thomas Scientific, New Jersey, USA), and successful amplicons were purified using the Geneaid PCR purification kit (Geneaid Biotech Ltd., Taipei, Taiwan). Sequencing was performed by a commercial company (Macrogen, Seoul, South Korea) on an automated Sanger sequencer.

**Table 2:**
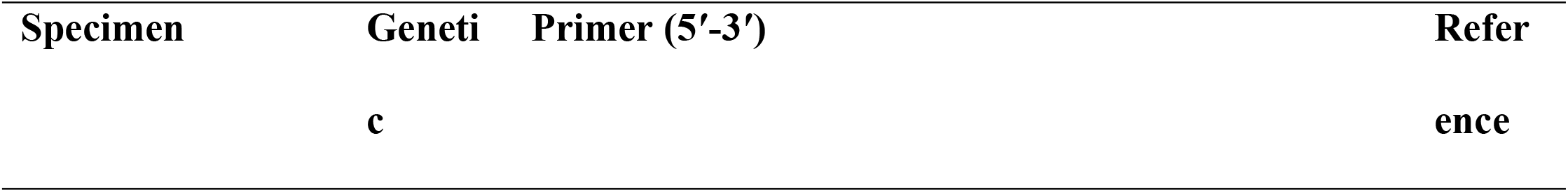
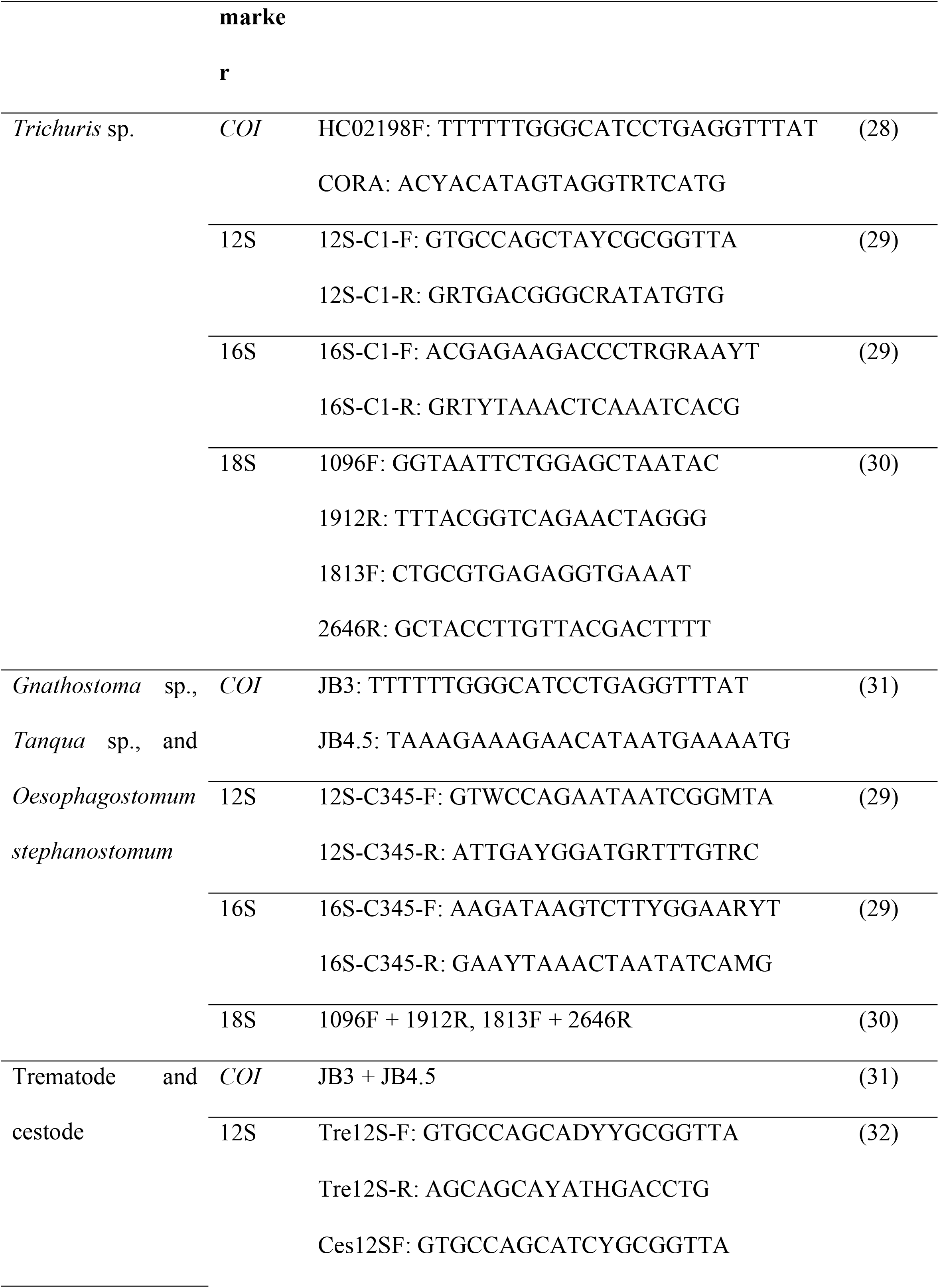
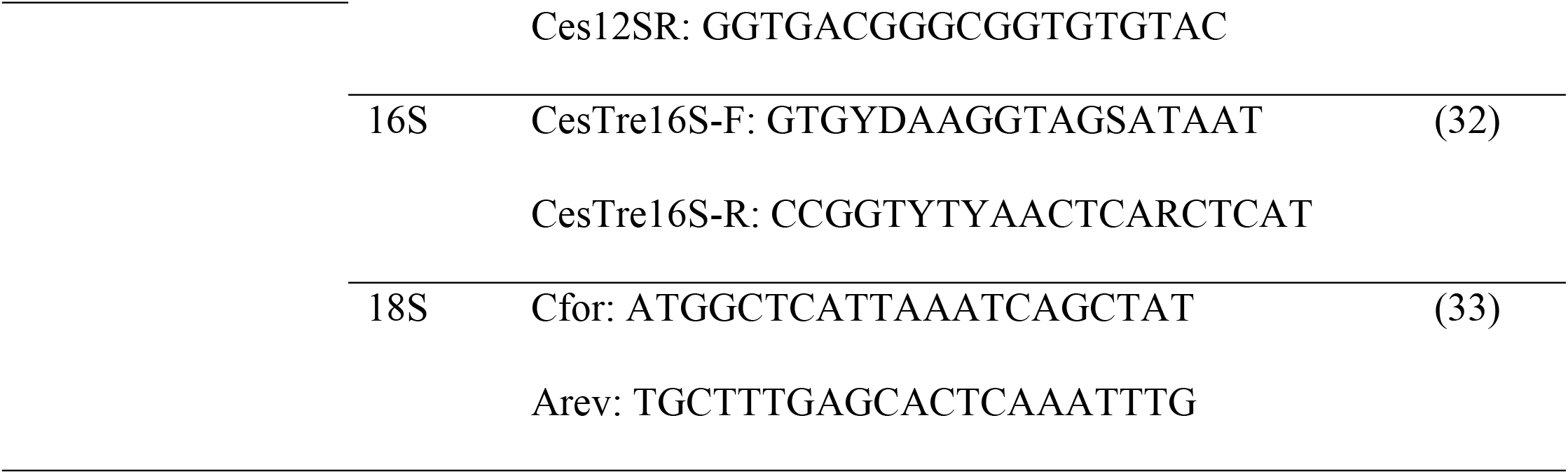
Primers used for ABIapp validation with actual specimens.

Electropherograms of the sequences were manually checked using Bioedit 7.0, and the sequences were input in NCBI BLAST to search for potentially similar species [24]. Multiple sequence alignments of species belonging to the same family/genus were then aligned together in ClustalX 2.1 [23]. The aligned sequences were checked in Bioedit 7.0 for ambiguous sites, and pairwise genetic distance (p-distance) was calculated using MEGA X [25]. The obtained genetic distances were then input into ABIapp to delimit each of the helminth specimens tested. The accuracy of ABIapp for species classification accuracy was determined based on the actual and predicted classification. The overall workflow for ABIapp validation is depicted in Fig 2.

**Fig 2.**
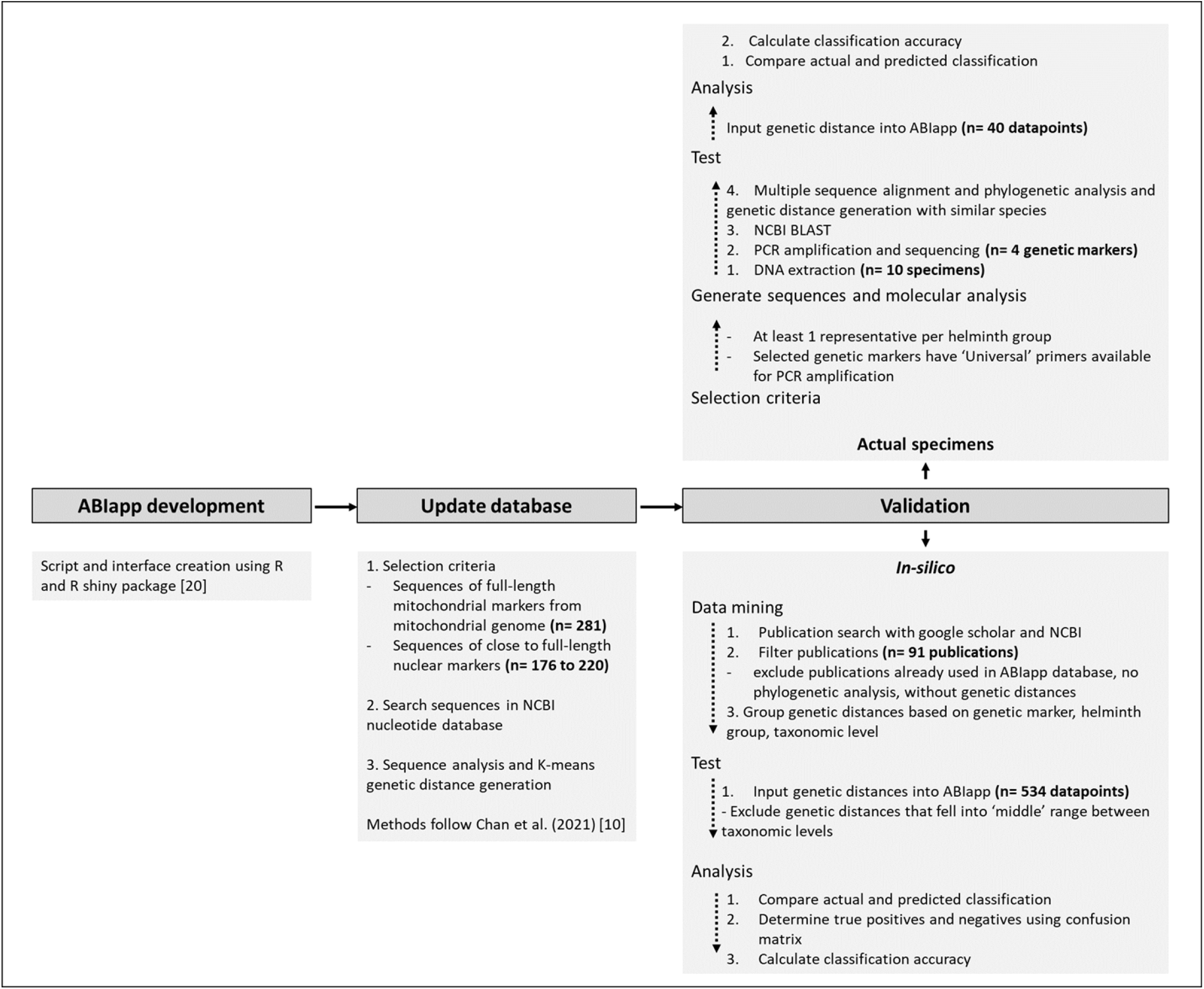
Workflow of ABIapp validation showing the three main steps – application development, update, and validation.

## Results and discussion

### In silico determination of the classification accuracy of ABIapp

Using 534 genetic distance values across the ten genetic markers for six helminth groups obtained from 91 publications (S3 Table), the classification accuracy of ABIapp was determined. Generally, an overall classification accuracy of 79% was achieved (ten genetic markers, six helminth groups, three taxonomic hierarchy levels). Figure 3 shows an intensity map of the classification accuracy of each genetic marker for each helminth group at their respective taxonomic levels, with the darker blue colors indicating higher accuracy. Comparing the classification accuracy per helminth group, nematode clade V was the highest at 93%, followed by trematode (Diplostomida) at 89% (S1A Fig). Among the three taxonomic hierarchy levels, there was no significant difference in the classification accuracy (76% to 80%) (S1B Fig). Comparing genetic markers, the average classification accuracy ranged from 57% to 91%, with the mitochondrial *COII* gene having the highest accuracy and the ITS1 region having the lowest (S1C Fig). Additionally, the mitochondrial genetic markers had better classification accuracy than the nuclear genetic markers, substantiating the validity and applicability of ABIapp for species delimitation using commonly applied mitochondrial genetic markers. A possible reason for the lower classification accuracy of the nuclear spacer regions could be the presence of repeat regions, thus complicating sequence alignment and genetic distance calculations across various publications [10, 34]. Also, as the nuclear 18S and 28S rRNA genes span a total of 4 and 12 domains, respectively, few publications utilize their complete length and difficulties could arise when different domains are used for genetic distance analysis [35].

**Fig 3.**
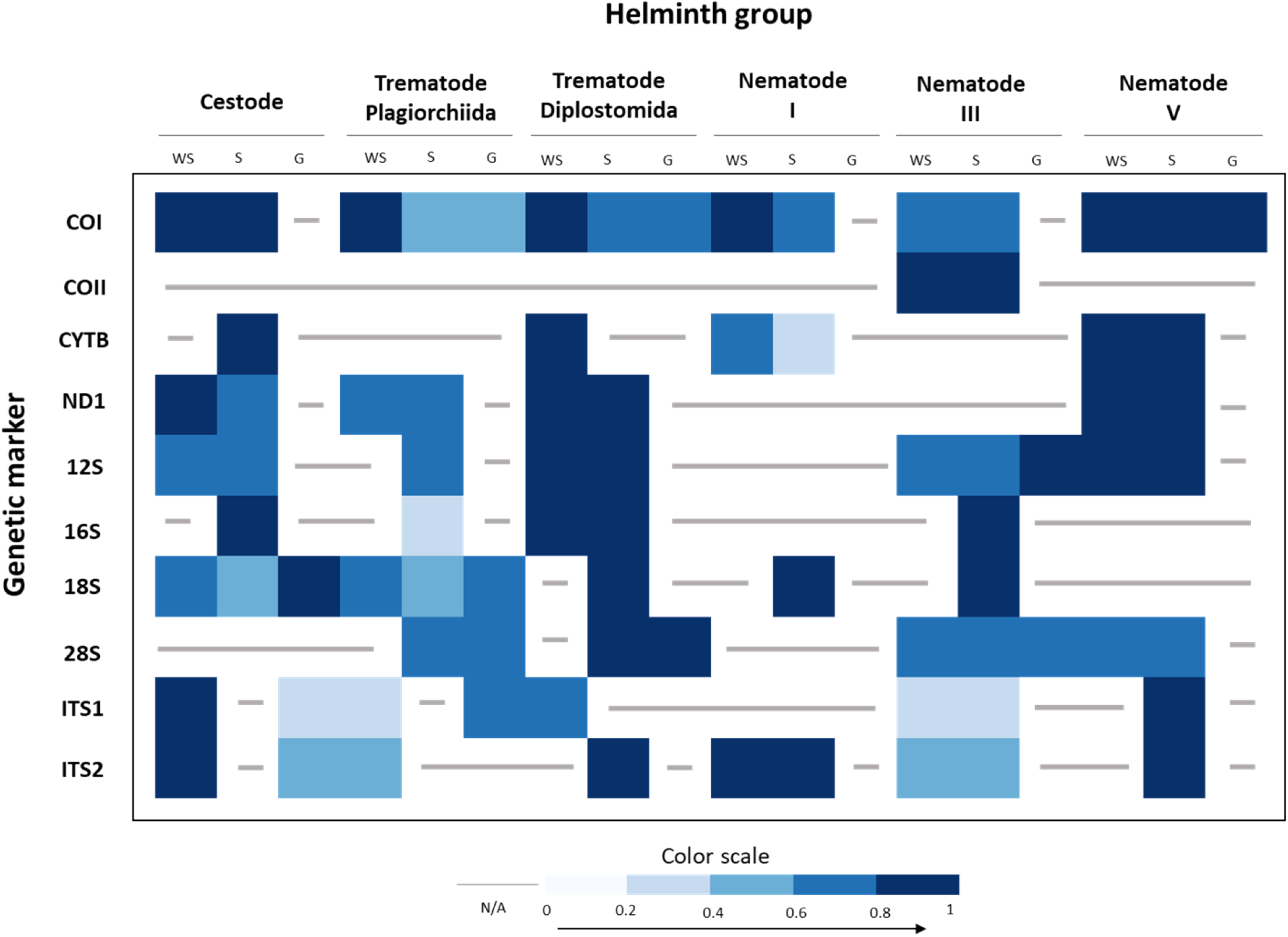
*In silico* species delimitation classification accuracy of ABIapp. The ten genetic markers tested are shown in the vertical column, while the six helminth groups are shown in the horizontal row. The taxonomic levels are represented by WS, S, and G, for within species, species, and genus, respectively. Classification accuracy percentage is represented by the intensity of the color scale, while the gray lines represent no data available (N/A).

Our results obtained through the *in silico* validation further support the robustness of ABIapp, with more than 75% accuracy achieved for eight out of ten genetic markers. Moreover, as the genetic distances tested were not restricted to groups or helminth hosts, ABIapp is applicable for a broad range of helminth species.

When we initially used the original dataset of cut-off genetic distances from Chan *et al*. [10], the overall classification accuracy through *in silico* validation was 69% (results not shown). Through increasing and updating the database of cut-off genetic distances, from approximately 91 to 281 helminth species in total (total number of species used for the mitochondrial genes), the overall classification accuracy was increased to 79%. We have therefore demonstrated the importance of increasing the number of species used to improve the classification accuracy of ABIapp. With the increasing trend of utilizing molecular genetic markers for helminth identification, the number of molecular sequences that are available in the NCBI database will undoubtedly increase [2]. Hence, the database of cut-off genetic distance values for ABIapp will be updated on a yearly basis, with the aim of improving the classification accuracy of ABIapp and broadening its applicability for users.

### Classification accuracy with actual specimens

From the ten helminth specimens tested with the mitochondrial *COI*, 12S, 16S, and nuclear 18S genes, the overall classification accuracy was 75%, with 30 out of 40 datapoints accurately classified. Figure 4 shows a representation of the classification accuracy of the ten helminth species tested. Of the six helminth groups, cestode and trematode (Plagiorchiida) had lower classification accuracy, while nematode clade I, nematode clade V, and trematode (Diplostomida) had 100% accuracy. These validation results are congruent with the *in silico* classification accuracy, where nematode clade V and trematode (Diplostomida) had the highest classification accuracy, while trematode (Plagiorchiida) had the lowest. Comparing the four genetic markers tested, the mitochondrial *COI* and 16S rRNA genes had the highest classification accuracy, at 90% (nine out of ten datapoints). This finding is especially important because the mitochondrial *COI* and 16S rRNA genes have proven to be useful genetic markers for molecular identification due to their high levels of sequence variation [29, 32, 36, 37].

**Fig 4.**
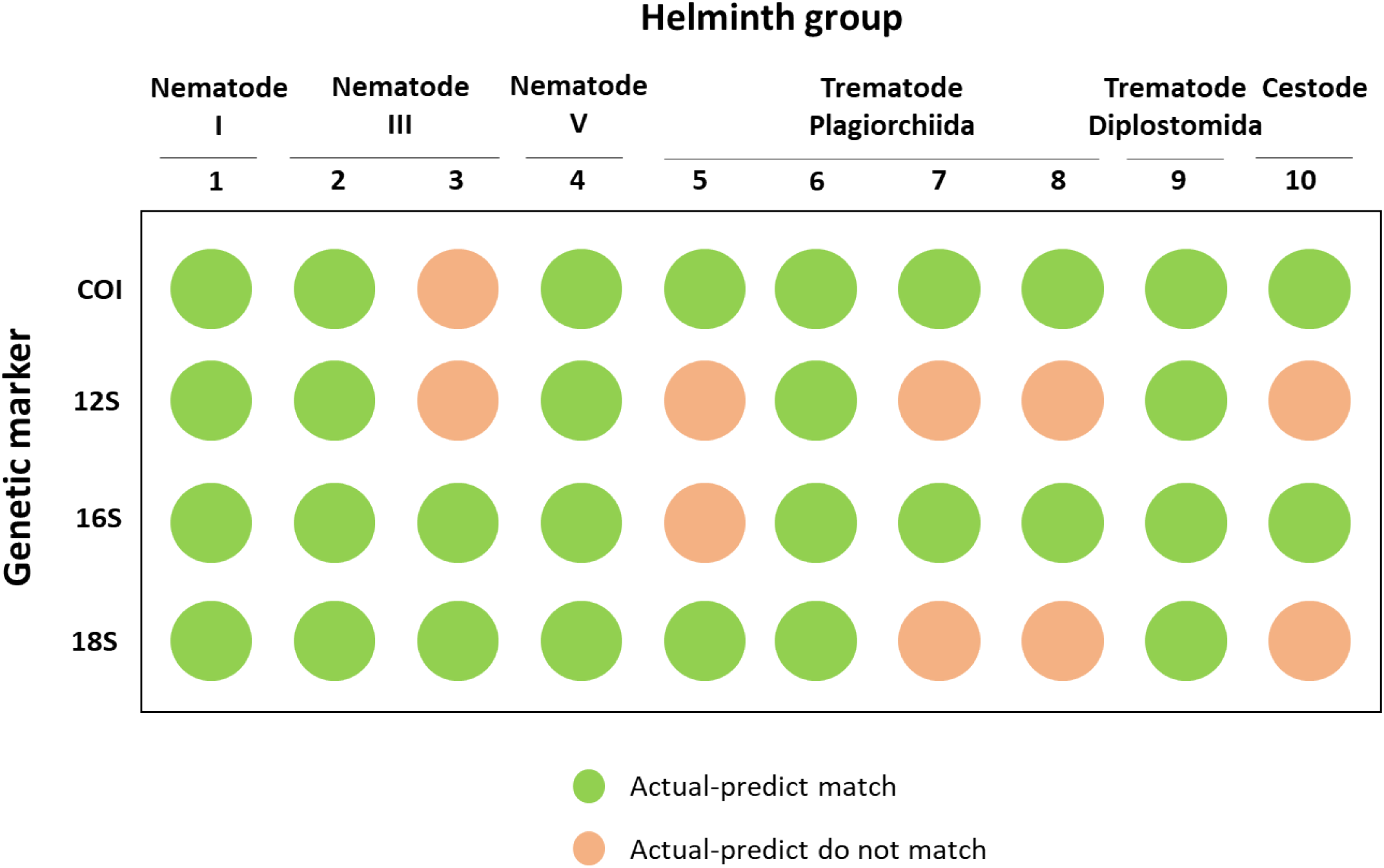
Classification accuracy of ABIapp using actual specimens. The four genetic markers tested are shown in the vertical column, while the ten helminth species are shown in the horizontal row (1-*Trichuris globulosa*, 2-*Gnathostoma* sp., 3-*Tanqua* sp., 4-*Oesophagostomum stephanostomum*, 5-*Gastrothylax* sp., 6-*Eurytrema* sp., 7-*Centrocestus formosanus*, 8-*Stellantchasmus falcatus*, 9-*Clinostomum* sp., 10-*Bertiella* sp.). Classification accuracy match is represented by the green- and pink-colored circles.

Incongruences between the actual and predicted results can be attributed to the proportion of sequences available in reference databases. For example, the *COI* gene has more reference sequences than the 12S rRNA gene, resulting in better classification accuracy when using the former in ABIapp. Another factor is the degree of sequence variation of the genetic marker. Using the 18S rRNA gene, although no sequence variation was observed (between our specimen of *S. falcatus* and the reference *S. falcatus*), ABIapp predicted that the two sequences were of different species (S5 Table). Inaccuracy in species delimitation using ABIapp occurred when the taxonomic boundaries for the species level using K-means ranged from 0 to 2.4%. Our findings also highlight the importance of using an additional genetic marker for species confirmation [38]. Another factor leading to incongruence can be the complicated taxonomic status for some groups of trematodes (e.g., within superfamily Opisthorchioidea) [32, 39, 40].

### Benefits of using ABIapp

ABIapp is a convenient tool for helminth species delimitation, as users can now easily determine whether their queried taxon is conspecific. In addition to its convenience for species delimitation, ABIapp offers multiple benefits to users.

First, with the estimated cut-off genetic distances incorporated into a webpage application and freely available as a package in R, ABIapp provides an accessible platform for users to utilize the application whenever they require. Second, ABIapp is useful as it is a straightforward application that only requires the use of genetic distance for helminth species delimitation or to determine a specimen’s taxonomic status. Compared with other more complex species delimitation methods, ABIapp requires only basic bioinformatics and molecular phylogenetic techniques to generate genetic distance values for analysis. Requiring just three types of input – genetic distance, helminth group, and genetic marker – ABIapp is easy to use, coupled with its user-friendly interface. Third, users need not be proficient in differentiating species based on morphology. With the declining pool of classical taxonomists in helminthology, reducing the need for expertise in morphological species identification is useful [41, 42]. Fourth, as the taxa selected for use in the genetic distance database cover many species of helminths found in various types of hosts and environments, the use of ABIapp is not restricted to a particular group of helminths. A broad user-base will undoubtedly aid in improving helminth species identification and delimitation, which will be beneficial for helminth taxonomy. Finally, by validating the classification accuracy of ABIapp through *in silico* methods and by using actual specimens, we have demonstrated that it is robust and applicable for a broad range of helminth species.

### Assumptions and limitations

ABIapp relies on certain assumptions and has some limitations that users should be aware of. First, as only genetic distances are input into the application, other information such as morphological characteristics or phylogenetic placements is not used to determine a specimen’s taxonomic status. Second, a species’ identity and taxonomic classification were assumed to be correct based on the information provided in the NCBI database. Third, genetic distances were obtained from other publications for the *in silico* validation; however, the various methods used in the studies to generate genetic distances were not accounted for. Finally, we did not consider the species complex status of some helminth species, although this can complicate species delimitation.

In conclusion, we have developed a convenient and user-friendly application to aid in helminth species delimitation that is applicable for a wide audience. The robustness of ABIapp for determining helminth taxonomic boundaries was also validated for its classification accuracy via *in silico* methods and the use of actual specimens. The database of genetic distances for ABIapp will be updated annually, to keep up to date with the increasing number of sequences available in molecular databases. ABIapp represents an accurate and validated application that is now readily available for researchers in the field of helminthology.

## Acknowledgments

We wish to acknowledge the Department of Helminthology, Faculty of Tropical Medicine, Mahidol University, Bangkok, for technical support and specimen collection.

## Supporting information files

S1A to S1E Tables: List of helminth species used in the ABIapp database

S2 Table: Genetic distances estimated using the K-means machine learning algorithm per helminth group input into ABIapp

S3 Table: List of publications used for the *in silico* validation of ABIapp

S4A to S4B Tables: *In silico* validation classification accuracy using ABIapp

S5 Table: ABIapp validation using actual specimens

S1A to S1C Figs: Average classification accuracy

## Credit author statement

Abigail Hui En Chan: Investigation, Methodology, Formal Analysis, Writing – Original Draft Preparation, Writing – Review & Editing

Urusa Thaenkham: Conceptualization, Methodology, Writing – Original Draft Preparation, Writing – Review & Editing

Tanaphum Wichaita: Methodology, Software, Writing – Review & Editing

Sompob Saralamba: Conceptualization, Methodology, Software, Writing – Original Draft Preparation, Writing – Review & Editing

